# Efficient recall of Omicron-reactive B cell memory after a third dose of SARS-CoV-2 mRNA vaccine

**DOI:** 10.1101/2022.02.20.481163

**Authors:** Rishi R. Goel, Mark M. Painter, Kendall A. Lundgreen, Sokratis A. Apostolidis, Amy E. Baxter, Josephine R. Giles, Divij Mathew, Ajinkya Pattekar, Arnold Reynaldi, David S. Khoury, Sigrid Gouma, Philip Hicks, Sarah Dysinger, Amanda Hicks, Harsh Sharma, Sarah Herring, Scott Korte, Wumesh KC, Derek A. Oldridge, Rachel I. Erickson, Madison E. Weirick, Christopher M. McAllister, Moses Awofolaju, Nicole Tanenbaum, Jeanette Dougherty, Sherea Long, Kurt D’Andrea, Jacob T. Hamilton, Maura McLaughlin, Justine C. Williams, Sharon Adamski, Oliva Kuthuru, Elizabeth M. Drapeau, Miles P. Davenport, Scott E. Hensley, Paul Bates, Allison R. Greenplate, E. John Wherry

## Abstract

Despite a clear role in protective immunity, the durability and quality of antibody and memory B cell responses induced by mRNA vaccination, particularly by a 3^rd^ dose of vaccine, remains unclear. Here, we examined antibody and memory B cell responses in a cohort of individuals sampled longitudinally for ∼9-10 months after the primary 2-dose mRNA vaccine series, as well as for ∼3 months after a 3^rd^ mRNA vaccine dose. Notably, antibody decay slowed significantly between 6- and 9-months post-primary vaccination, essentially stabilizing at the time of the 3^rd^ dose. Antibody quality also continued to improve for at least 9 months after primary 2-dose vaccination. Spike- and RBD-specific memory B cells were stable through 9 months post-vaccination with no evidence of decline over time, and ∼40-50% of RBD-specific memory B cells were capable of simultaneously recognizing the Alpha, Beta, Delta, and Omicron variants. Omicron-binding memory B cells induced by the first 2 doses of mRNA vaccine were boosted significantly by a 3rd dose and the magnitude of this boosting was similar to memory B cells specific for other variants. Pre-3^rd^ dose memory B cell frequencies correlated with the increase in neutralizing antibody titers after the 3^rd^ dose. In contrast, pre-3^rd^ dose antibody titers inversely correlated with the fold-change of antibody boosting, suggesting that high levels of circulating antibodies may limit reactivation of immunological memory and constrain further antibody boosting by mRNA vaccines. These data provide a deeper understanding of how the quantity and quality of antibody and memory B cell responses change over time and number of antigen exposures. These data also provide insight into potential immune dynamics following recall responses to additional vaccine doses or post-vaccination infections.

**Graphical Summary:** 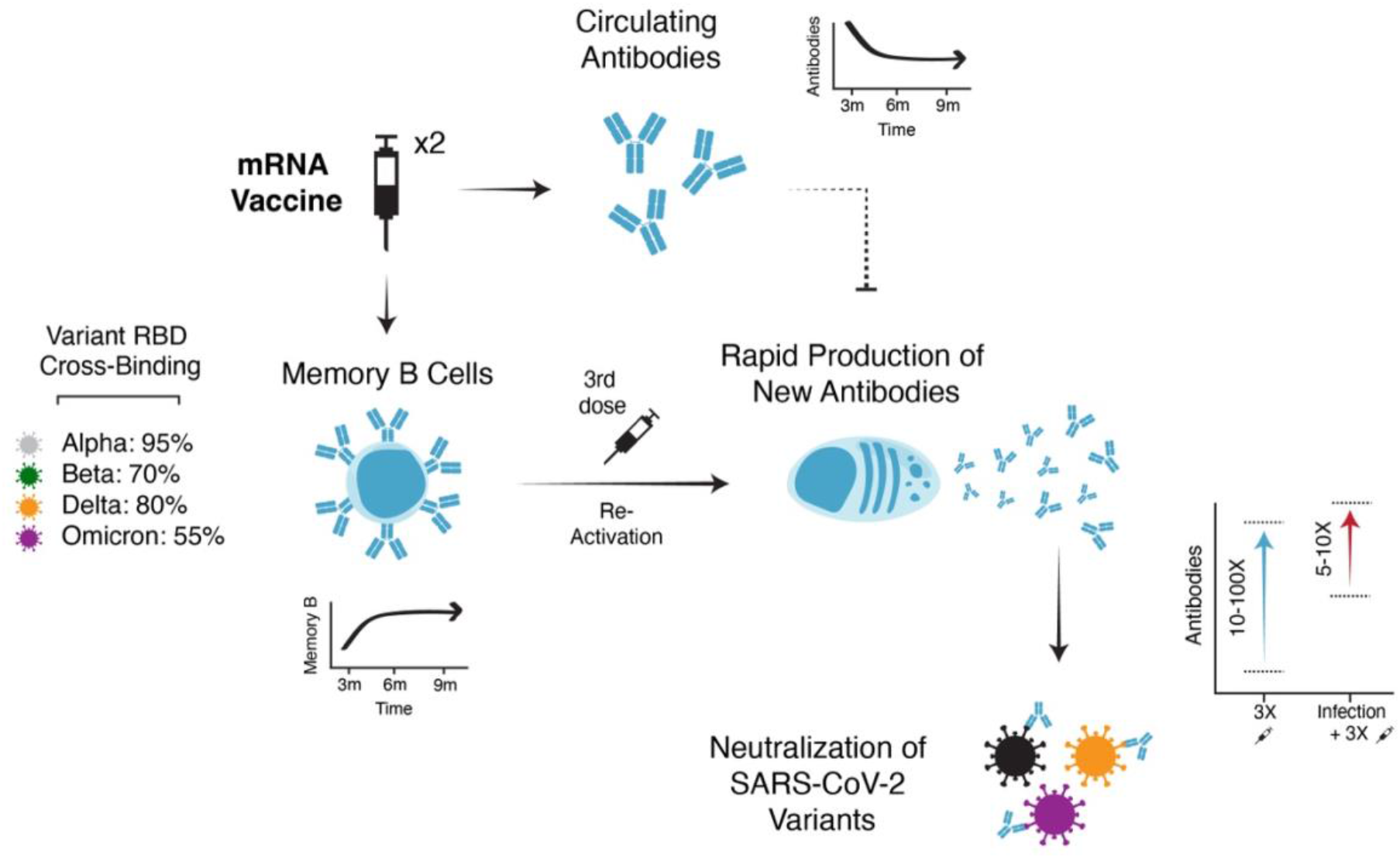

## Introduction

SARS-CoV-2 infections continue to cause significant morbidity and mortality worldwide (*1*). Since the virus was identified in late 2019, several SARS-CoV-2 variants of concern (VOC) have emerged. Mutations found in SARS-CoV-2 variants, particularly those in the Spike glycoprotein, can alter viral transmission and immune recognition (*2–4*). Of these VOC, the Delta (B.1.617.2) variant had considerable impact due to its increased infectivity and partial escape from neutralizing antibodies (*5, 6*). Most recently, scientists in South Africa identified and characterized the Omicron (B.1.1.529) variant (*7*). In the weeks following identification, Omicron spread rapidly, outcompeting Delta to become the dominant variant in the US and many parts of the world.

A major concern about Omicron is the large number of mutations in the Spike protein, including ∼15 amino acid changes in the Spike receptor binding domain (RBD). *In vitro* data indicate that these mutations have a substantial effect on evading antibody responses in convalescent or mRNA vaccinated (Pfizer BNT162b2 or Moderna mRNA-1273) individuals. This effect is more pronounced than other VOC, with a ∼10 to ∼40-fold reduction in neutralization capacity compared to wild-type virus using either pseudovirus or live virus neutralization assays, and little to no neutralizing activity against Omicron detected at >6 months after the primary 2-dose vaccine series (*8–11*).

In addition to circulating antibodies, memory B cells represent an important source of long-term immunity (*12, 13*). In contrast to antibodies that decline over the first 3-6 months post vaccination (*14*), antigen-specific memory B cells appear highly stable over time (*15*). Upon re-exposure to antigen, either through vaccination or infection, these memory B cells can differentiate into antibody secreting cells and rapidly produce new antibodies (*16*). Indeed, recent non-human primate studies of mRNA vaccination highlight recall antibody responses from memory B cells as a key factor in protection from severe COVID-19 pathology in the lungs (*17*). Previous work has shown that mRNA vaccines induce robust memory B cell responses that continue to evolve via germinal centers for months after primary vaccination (*15, 18–21*). As a result, immunization with mRNA vaccines encoding the original Wuhan Spike protein generates a population of high-affinity memory B cells that can bind the Alpha, Beta, and Delta variants and produce neutralizing antibodies upon restimulation.

Serologic data indicate that antibody responses to Omicron can be at least partially boosted in the short-term (up to ∼1 month) after a 3^rd^ vaccine dose (*22–25*), suggesting that immunological memory generated by 2-dose vaccination has some reactivity against the Omicron Spike protein. A 3^rd^ vaccine dose also provides increased protection from Omicron variant infection (*26*). However, it is unclear how long these boosted antibody responses to Omicron may last and what percent of memory B cells retain binding to Omicron and other variants. Moreover, the dynamics of memory B cell responses in humans are poorly understood, and whether boosting with the original Wuhan Spike can overcome antigenic changes by efficiently reactivating Omicron-binding memory B cells is unknown. Finally, it remains unclear what features of immunity induced by 2-dose vaccination determine optimal boosting following a 3^rd^ vaccine dose, and how immune responses are affected by additional antigen encounters beyond a 3-dose vaccine schedule. The answers to these questions should inform how to optimize the use of additional vaccine doses for protection against Omicron and future VOC.

## Results

### Study Design

We examined antibody and memory B cell responses to SARS-CoV-2 in a longitudinal cohort of 61 individuals receiving mRNA vaccines (Pfizer BNT162b2 or Moderna mRNA-1273). This cohort has been previously described through 6 months post-2 doses of mRNA vaccine (*15, 18, 27*). 45 individuals were infection naïve and 16 had recovered from a prior infection. Paired serum and peripheral blood mononuclear cell (PBMC) samples were collected at 10 different timepoints, ranging from pre-vaccine baseline through ∼9-10 months post-primary 2-dose vaccination, as well as prior to a 3^rd^ vaccine dose, ∼ 2 weeks post-3^rd^ dose, and ∼3 months post-3^rd^ dose (figure 1A). Seven individuals had a confirmed post-vaccination (i.e., breakthrough) infection during the study period and are indicated in all analyses. Additional cohort information is provided in table S1.

**Figure 1.**
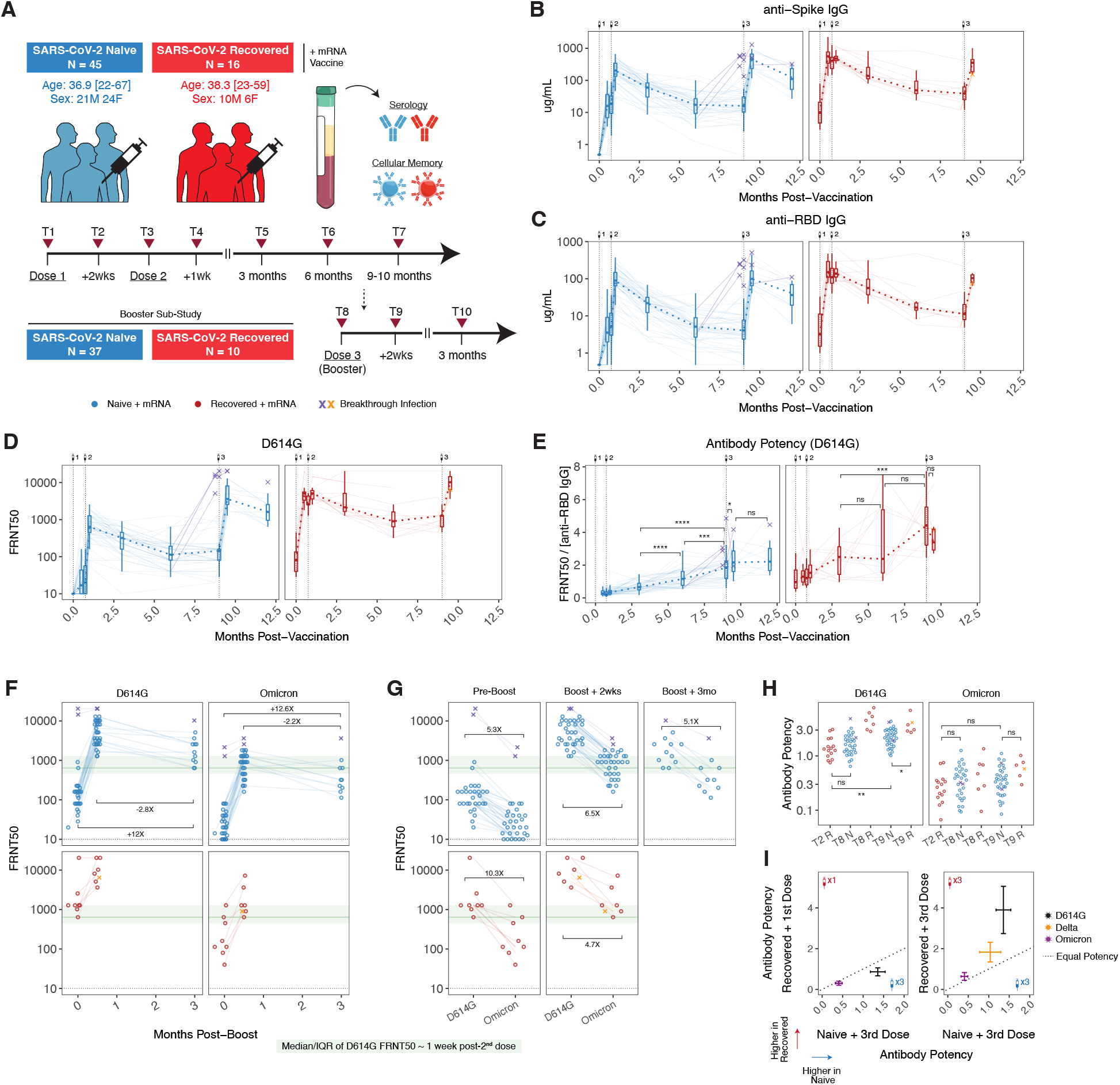
Antibody responses after 2 and 3 doses of mRNA vaccine. **A)** Study design and cohort characteristics. **B)** anti-Spike and **C)** anti-RBD IgG concentrations over time in plasma samples from vaccinated individuals. **D)** Pseudovirus (PSV) neutralization titers against wild-type D614G Spike protein over time in plasma samples from vaccinated individuals. Data are represented as focus reduction neutralization titer 50% (FRNT50) values. **E)** Antibody neutralization potency against D614G over time. Potency was calculated as neutralizing titer (FRNT50) divided by the paired concentration of anti-RBD IgG. **F and G)** Plasma neutralizing activity against D614G and Omicron before and after booster vaccination. Dotted lines indicate limit of detection for the assay. Green boxes and lines indicate interquartile range (IQR) and median of D614G neutralizing titers ∼1 week following the second vaccine dose in SARS-CoV-2 naïve subjects. **H and I)** Comparison of antibody potency against D614G, Delta, and Omicron between SARS-CoV-2 naïve and previously infected vaccinees. For **I**, bars indicate mean with 95% confidence intervals. Statistics were calculated using unpaired non-parametric Wilcoxon test with BH correction for multiple comparisons. *P < 0.05; **P < 0.01; ***P < 0.001; ****P < 0.0001; ns, not significant.

### Antibody Responses

As previously described, 2-dose mRNA vaccination in previously uninfected individuals induced high titers of anti-Spike and anti-RBD binding antibodies, as well as neutralization titers against wild-type (D614G) pseudovirus (figure 1B-D) (*15, 18*). Vaccination in individuals with a prior SARS-CoV-2 infection (commonly referred to as “hybrid immunity”) resulted in even higher antibody titers after the first dose, consistent with an anamnestic response from pre-vaccination immunological memory (figure 1B-D). Although the absolute magnitude of antibody responses differed between COVID naïve and COVID recovered vaccinees, the decay rates from peak responses (∼1 week after the 2^nd^ dose) to ∼6 months post-primary vaccination were similar in both groups (figure 1B-D). This decay slowed over time and antibody titers stabilized between 6- and 9-months post-vaccination for both groups, with little to no decrease in neutralizing antibody titers after 6 months (figure 1B-D). These findings are consistent with ongoing antibody production from long-lived plasma cells in the later phases of immune memory after vaccination.

To evaluate the quality of antibody responses, we calculated an antibody potency index based on the ratio of neutralization titers to the total concentration of anti-RBD binding IgG. Antibody potency increased significantly over time after the 2^nd^ vaccine dose, with a continued increase in potency from 6 to 9 months post-vaccination in the infection naïve group as antibody concentrations began to plateau (figure 1E). These observations suggest decay of lower quality antibody from short-lived antibody secreting cells, as well as continued emergence of post-germinal center affinity matured plasma cells over time that produce higher quality antibody later in the response. This improvement in the quality of antibody for at least 9 months is also consistent with a recent report demonstrating the continued presence of Spike-binding germinal center B cells in axillary lymph nodes at 29 weeks post-vaccination (*20*).

In addition to the primary 2-dose vaccine series, most of our cohort went on to receive a 3^rd^ dose of mRNA vaccine. A 3^rd^ dose of vaccine in infection naïve individuals significantly increased binding and neutralizing antibodies, with both reaching a similar level to that observed in previously infected individuals after the 2-dose vaccine series (figure 1B-D). A 3^rd^ dose of mRNA vaccine in these COVID recovered individuals (i.e. a 4^th^ exposure) also significantly boosted antibody responses; however, the relative magnitude of this increase was less than observed in the recall response after the initial 2-dose vaccine series (figure 1B-D). Several individuals in this cohort also experienced breakthrough infections after 2 or 3 doses of vaccine. Although the sample size was limited (N=3), infection following 2-dose vaccination appeared to boost antibodies to similar levels as previous infection with 2 doses of vaccine (figure 1B-D), suggesting that the total number of antigen exposures may be as important as the relative order of exposure to infection and vaccination.

To quantify neutralizing capacity of vaccine-induced antibody responses against VOCs, we generated pseudotyped viruses encoding the Delta and Omicron Spike proteins. Consistent with previous reports, neutralizing titers against Omicron were significantly reduced relative to D614G, with ∼20% of individuals having neutralization titers below limit of detection at ∼9 months post-primary vaccination (figure 1F-G). Following a 3^rd^ dose, neutralizing titers to Omicron increased by ∼30-fold, with similar kinetics and magnitude of boost as neutralizing antibodies against D614G (figure 1F-G). Although neutralization against Omicron declined 2.2-fold from peak levels between 2 weeks post 3^rd^ dose and 3 months post 3^rd^ dose, titers remained 12.6-fold above pre-3^rd^ dose baseline in COVID-naïve individuals (figure 1F), indicating that an additional vaccine dose has a lasting benefit for antibodies against Omicron. In paired comparisons for individuals, Omicron neutralizing antibodies were on average 6.5-fold lower than neutralizing titers to D614G at the peak response after the 3^rd^ vaccine dose and 5.1-fold lower 3 months later. Despite the relative loss of neutralizing activity, peak Omicron neutralizing titers after the 3^rd^ dose were comparable to neutralizing titers against D614G ∼1 week after the 2^nd^ dose, where clinical efficacy has previously been defined (figure 1F-G) (*28*). Moreover, Omicron neutralizing titers at 3 months post-3^rd^ dose were similar to or higher than pre-3^rd^ dose D614G neutralizing titers (figure 1F-G). Finally, to investigate potential differences in the quality of antibody recall responses, we compared antibody potency 2 weeks after the first dose of mRNA vaccine in individuals with pre-existing immune memory from infection to antibody potency 2 weeks after a 3^rd^ dose of mRNA vaccine in previously uninfected individuals. Previous reports have suggested that infection generates greater antibody potency and breadth than 2-dose mRNA vaccination alone (*29*). In this cohort, however, antibody potency against both D614G and Omicron was slightly higher following a 3^rd^ dose of mRNA vaccine compared to recall responses following the 1^st^ dose in SARS-CoV-2 recovered individuals (figure 1H-I), suggesting that a 3^rd^ dose of mRNA drives antibody potency to similar levels as “hybrid immunity”. Of note, potency did continue to increase in SARS-CoV-2 recovered individuals between the initial recall response to vaccine and 9+ months post-vaccination (figure 1E,H-I). This finding indicates there may be ongoing evolution after a recall response that can result in further improvement of antibody potency. Taken together, these data demonstrate that antibody responses, including neutralizing antibodies to Omicron, are effectively boosted by a 3^rd^ dose of mRNA vaccine with sustained benefit at ∼3 months post-3^rd^ dose.

### Memory B Cell Responses

We next investigated B cell responses to mRNA vaccination. Antigen-specific B cell responses were quantified from bulk PBMCs by flow cytometry using fluorescently labelled SARS-CoV-2 Spike and RBD probes as previously described(*15, 18, 30*). Influenza Hemagglutinin (HA) was used as a historical antigen for a specificity control. Plasmablasts were identified as CD20-CD38++ non-naïve B cells. Memory B cells were identified as CD20+ CD38lo/int non-naïve B cells. Full gating strategies are shown in figure S1.

Consistent with our plasma antibody data, re-exposure to SARS-CoV-2 antigen, either through a 3^rd^ mRNA vaccine dose or breakthrough infection, resulted in significant expansion of Spike-binding plasmablasts ∼1 week after antigen encounter (figure 2A-B). Overall, the rapid emergence of antibody-secreting cells following antigen re-encounter is consistent with recall from a pool of memory B cells. These data are consistent with findings after viral challenge in SARS-CoV-2 mRNA-vaccinated monkeys, where anamnestic antibody responses from memory B cells were identified as a major protective mechanism (*17*).

**Figure 2.**
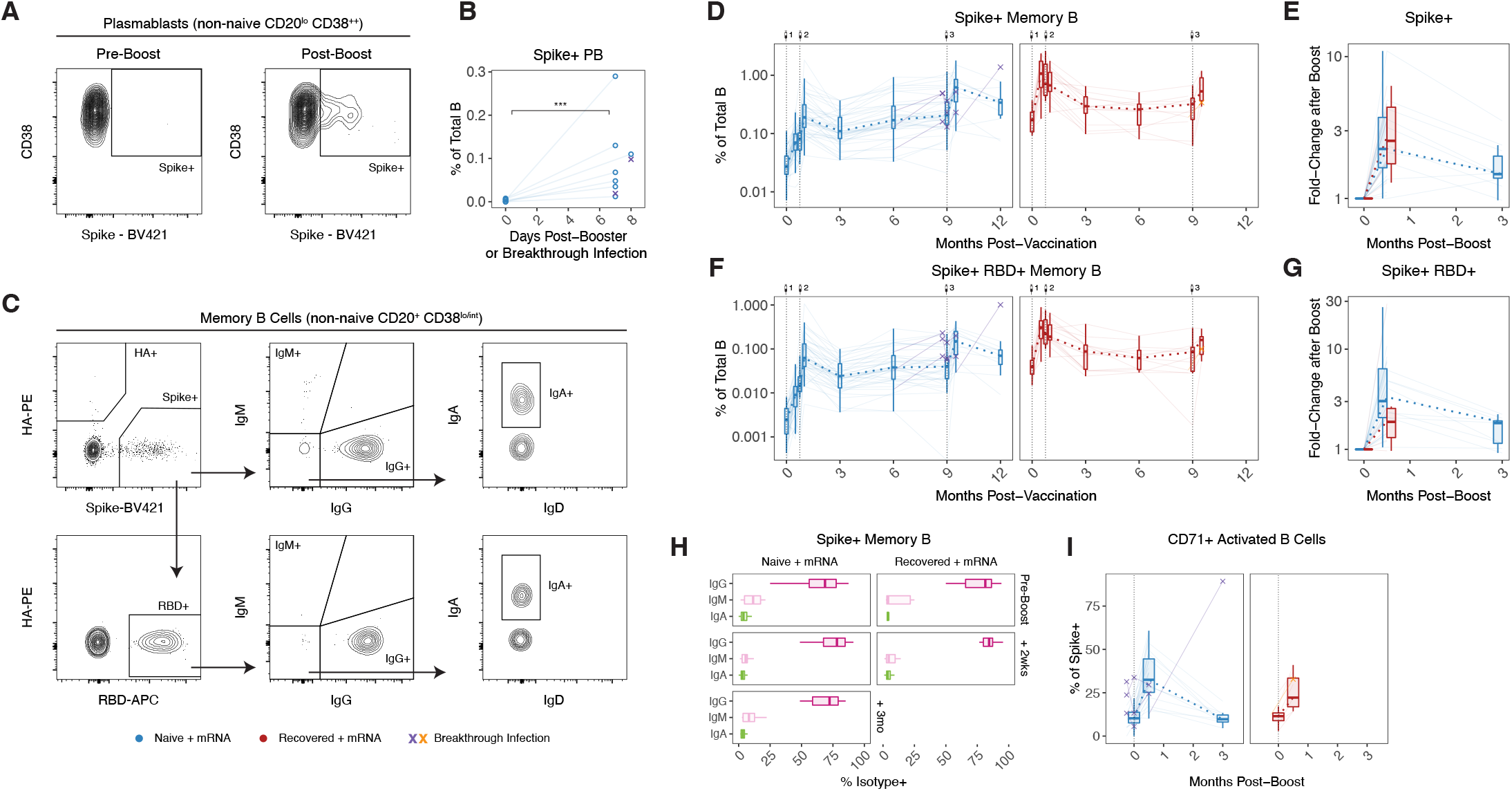
Memory B cell responses after 2 and 3 doses of mRNA vaccine. **A)** Flow cytometry gating strategy for SARS-CoV-2-specific plasmablasts. **B)** Frequency of Spike+ plasmablasts ∼1 week after booster vaccination or post-vaccine breakthrough infection. Data are represented as a percentage of total B cells. **C)** Flow cytometry gating strategy for SARS-CoV-2-specific memory B cells. **D and E)** Frequency of Spike+ and **F and G)** Spike+ RBD+ memory B cells over time in PBMCs from vaccinated individuals. Data are represented as a percentage of total B cells. **H)** Isotype composition of Spike+ memory B cells in vaccinated individuals pre- and post-boost. **I)** Activation status of Spike+ memory B cells over time in vaccinated individuals following booster vaccination. Statistics were calculated using unpaired non-parametric Wilcoxon test. *P < 0.05; **P < 0.01; ***P < 0.001; ****P < 0.0001; ns, not significant.

We previously demonstrated that mRNA vaccines induce durable and functional memory B cells to SARS-CoV-2 that are stable for at least 6 months after vaccination (*15*). Here, we extended these observations by tracking responses further into the memory phase. Spike-and RBD-specific memory B cell numbers continued to remain highly stable through at least 9 months post-vaccination in both SARS-CoV-2 naïve and previously infected individuals with no evidence of decline in numbers from 6 to 9 months post-primary vaccination (figure 2C-E). Notably, 34/35 SARS-CoV-2 naïve individuals had Spike- and RBD-specific memory B cell frequencies above their pre-vaccine baseline at the 9-month timepoint, highlighting continued the durability of mRNA-vaccine induced cellular immunity.

Upon receipt of a 3^rd^ mRNA vaccine dose, these memory B cells expanded in number. At ∼2 weeks post-boost there was a 2.2-fold fold increase in Spike-specific and a 3.3-fold increase in RBD-specific memory B cells in infection naïve individuals, with similar boosting for Spike- and RBD-specific memory B cells in COVID recovered vaccinees (figure 2D-G). By 3 months post 3^rd^ vaccine dose in infection naïve subjects, memory B cells had declined from peak levels but still remained ∼1.5-fold more abundant than before boosting (figure 2D-G). The majority of memory B cells after 2 doses of mRNA vaccine were IgG+ with a small percentage of IgM+ and IgA+ cells (figure 2H). A 3^rd^ vaccination did not markedly change the isotype composition of the memory response (figure 2H). A 3^rd^ dose of vaccine also induced a population of CD71+ activated B cells, consistent with reactivation of memory B cells (figure 2I). This activation status, however, transitioned back to a resting memory phenotype by 3 months post-3^rd^ dose. Thus, memory B cells are rapidly reactivated by re-exposure to antigen, either through infection or vaccination, and this reactivation is associated with induction of plasmablasts, numerical expansion of memory B cells, and re-establishment of B cell memory.

### Omicron- and Other Variant-Specific Memory B Cell Responses

A major question is whether vaccination with the original Wuhan Spike protein sequence induces effective immunological memory to VOCs including Omicron, and if so, whether Omicron-specific memory B cells can be efficiently boosted by subsequently re-vaccinating with wild-type vaccine. To investigate if mRNA vaccine-induced memory B cells were capable of recognizing the Omicron variant and determine how Omicron binding related to specificity for other VOCs, we designed a modified flow cytometry panel with antigen probes for 9 SARS-CoV-2 antigens. This panel included full-length Spike, N-terminal domain (NTD), S2 domain, wild-type (WT, Wuhan-Hu-1) RBD, and 4 variant RBDs (Alpha, Beta, Delta and Omicron). Nucleocapsid (N) was included for a response indicative of infection but not vaccination. For this analysis, B cells were enriched from total PBMCs by negative selection at 3 timepoints: pre-3^rd^ dose, ∼2 weeks after the 3^rd^ dose, and ∼3 months the 3^rd^ dose. Representative plots and gating strategy are shown in figure 3A-B, and figure S2. Briefly, antigen-specificity and phenotype were identified according to the following gating strategy: Spike+ memory B cells were first identified from total memory B cells as described above. Spike+ memory B cells were subsequently gated based on co-binding to NTD or S2 probes. Memory B cells that were Spike+ but did not bind NTD or S2 were then examined for binding to the WT RBD probe, as well as variant RBD probes.

**Figure 3.**
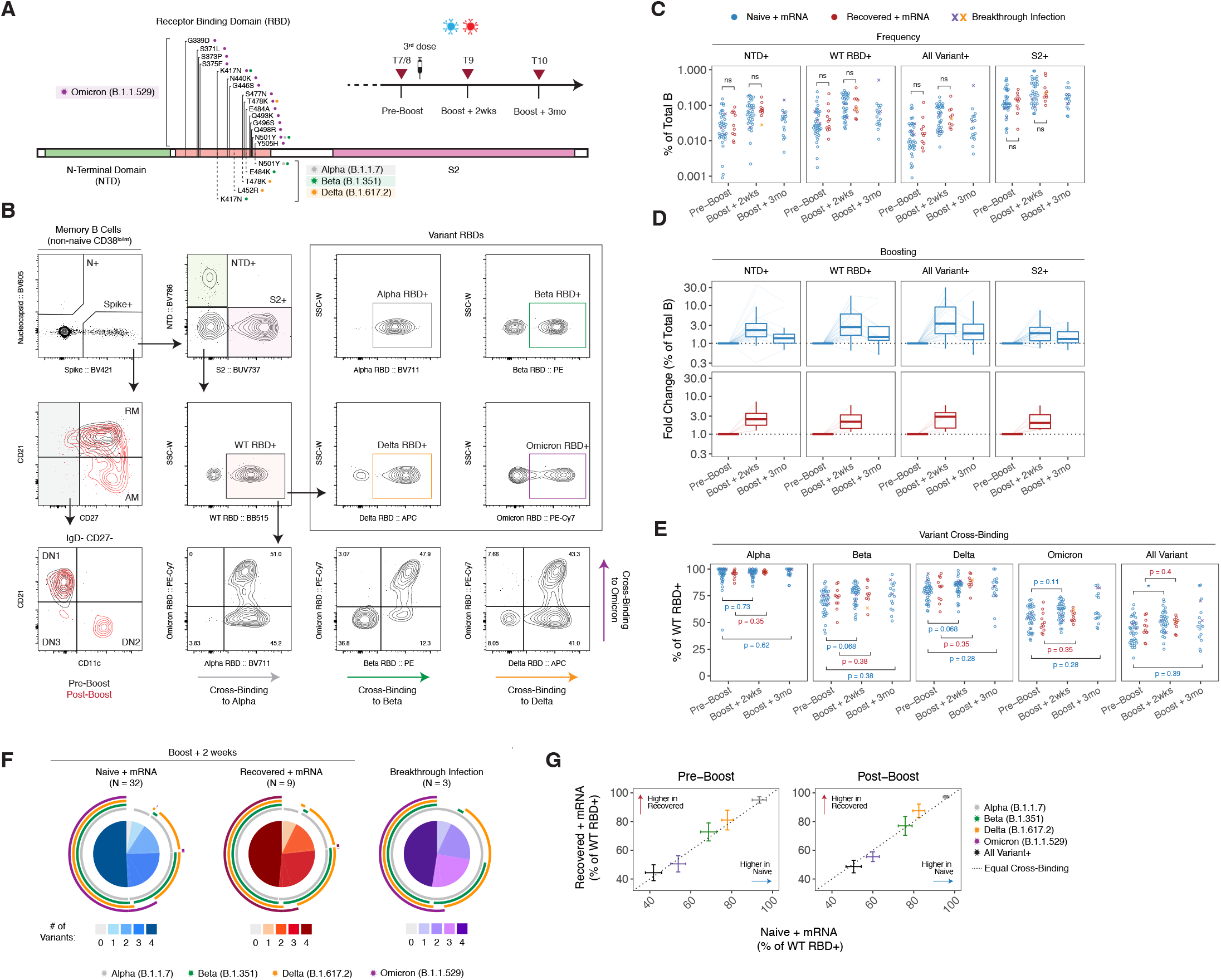
Variant-reactive memory B cell responses after 2 and 3 doses of mRNA vaccine. **A)** Experimental design and **B)** flow cytometry gating strategy for SARS-CoV-2 variant-reactive memory B cells. **C)** Frequency of NTD+, WT RBD+, All Variant RBD+, and S2+ memory B cells in vaccinated individuals pre- and post-boost. **D)** Fold change in the frequency of antigen-specific memory B cells after booster vaccination relative to paired pre-boost samples. Dotted lines indicate no change in frequency compared to pre-boost samples. **E)** Variant cross-binding of RBD-specific memory B cells in vaccinated individuals. Data are represented as a percentage of WT RBD+ cells. **F)** Boolean analysis of variant cross-binding memory B cell populations in vaccinated individuals ∼2 weeks after 3^rd^ vaccination or at a cross-sectional timepoint in individuals with a post-vaccine breakthrough infection. Pie charts indicate the fraction of WT RBD+ memory B cells that cross-bind zero, one, two, three, or four variant RBDs. Colored arcs indicate cross-binding to specific variants. **G)** Comparison of RBD variant cross-binding between SARS-CoV-2 naïve and previously infected vaccinees before and ∼2 weeks after 3rd vaccination. For **G**, bars indicate mean with 95% confidence intervals. Statistics were calculated using unpaired non-parametric Wilcoxon test with BH correction for multiple comparisons. *P < 0.05; **P < 0.01; ***P < 0.001; ****P < 0.0001; ns, not significant.

As previously described, 2-dose vaccination induced memory B cells specific for all domains of the Spike protein, with S2 representing the immunodominant part of the response (*15*). A 3^rd^ dose of mRNA vaccine boosted NTD-, WT RBD- and S2-specific memory B cells with a 2.4-fold increase in NTD-specific memory B cells, 3.2-fold increase in WT RBD-specific memory B cells, and a 1.9-fold increase in S2-specific memory B cells (figure 3C-D). Notably, memory B cells that recognized all RBD variants (Alpha, Beta, Delta, and Omicron) simultaneously had the greatest fold change after the boost (3.8-fold; figure 3C-D), indicating that a 3^rd^ exposure to wild-type Spike was sufficient to expand memory B cells targeting multiple different VOCs.

To further investigate variant-specificity within the memory B cell compartment, we quantified cross-binding to different VOC probes as a percentage of WT RBD-binding cells. 9 months after primary vaccination, >90% of memory B cells that bound WT RBD also bound Alpha RBD containing a single N501Y substitution (figure 3E). The L452R and T478K mutations found in Delta resulted in moderate loss of binding, with ∼80% of WT RBD+ cells still able to cross-bind Delta RBD (figure 3E). The K417N, E484K, and N501 mutations found in Beta were more immune evasive than Delta, with ∼70% of WT RBD+ memory B cells able to cross-bind Beta RBD (figure 3E). Notably, ∼55% of WT RBD-binding memory B cells after 2 doses of vaccine were still able to cross-bind Omicron RBD (figure 3E). Most Omicron RBD-specific memory B cells were also capable of recognizing Alpha, Beta and Delta RBDs, though binding overlap was less complete for Omicron and Delta compared to other combinations (figure 3F), likely due to the L452R mutation that is found in Delta but not Omicron. As a result, a considerable fraction (∼40-50%) of RBD-specific memory B cells could bind all 4 VOC RBDs simultaneously (figure 3E-F). These “All Variant+” memory B cells may represent a source of broad protection that is resilient to future VOCs. Boosting appeared to slightly increase memory B cell cross-binding to Beta, Delta, and Omicron RBDs (figure 3E), and the overall VOC binding profiles were similar in both COVID-naïve and prior COVID vaccinees (figure 3G).

To investigate how memory B cells with different antigen-specificities were recruited by a 3^rd^ dose of vaccine containing Wuhan Spike, we examined the activation phenotype of these cells ∼2 weeks after the 3^rd^ vaccine dose. Memory B cell activation state was defined based on expression of CD21, CD27, and CD11c (figure 3B). CD21-CD27+ B cells were identified as activated memory (AM) and CD21+ CD27+ cells were identified as resting memory (RM). CD27-cells were split into double negative (DN) 1, 2, and 3 subsets based on CD21 and CD11c staining. As expected, re-vaccination with a 3^rd^ dose induced a clear transition from a resting memory to an activated memory B cell phenotype in Spike-specific cells (figure 4A). DN1, DN2, and DN3 cells only represented a small fraction of the overall response. NTD+, RBD+, and S2+ memory B cell activation states clustered together (figure 4A), indicating that antigen re-exposure activated memory B cells specific for all parts of the Spike protein.

**Figure 4.**
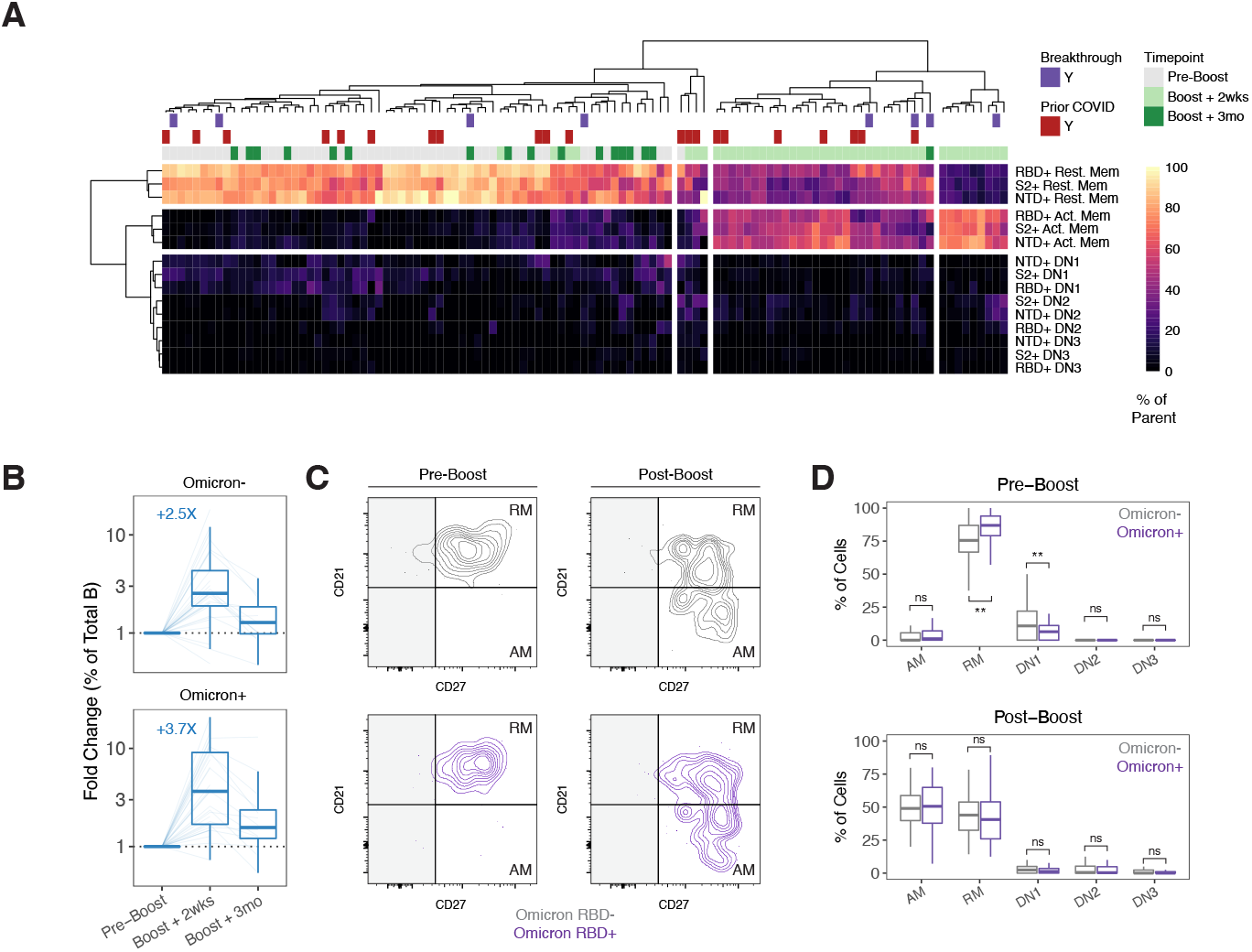
Activation of Omicron-specific B cell memory after a 3^rd^ dose of mRNA vaccine. **A)** Heatmap and hierarchal clustering of memory B cell activation status by antigen specificity at pre- and post-3^rd^ dose timepoints. Prior COVID infection and/or post-vaccine breakthrough infection are indicated. **B)** Fold change in the frequency of Omicron RBD-binding versus non-binding memory B cells after booster vaccination relative to paired pre-boost samples. Dotted lines indicate no change in frequency compared to pre-boost samples. **C)** Representative flow cytometry plots for activation phenotype of Omicron RBD-binding versus non-binding memory B cells. **D)** Frequency of activated memory (AM), resting memory (RM), or double negative (DN) subsets in Omicron RBD-binding versus non-binding memory B cells before and ∼2 weeks after 3rd vaccination. For **B-D**, analysis was restricted to SARS-CoV-2 naïve vaccinees. Statistics were calculated using paired non-parametric Wilcoxon test with BH correction for multiple comparisons. *P < 0.05; **P < 0.01; ***P < 0.001; ****P < 0.0001; ns, not significant.

Within the RBD-specific memory B cell population, it has been unclear if variant-specific memory B cells could be activated by a 3^rd^ dose of vaccine, and if so, would there be a preference for VOC non-binders over VOC binders. Here, a 3^rd^ dose of mRNA vaccine encoding the original Wuhan Spike boosted Omicron RBD-binding memory B cells by 3.7-fold compared to 2.5-fold for memory B cells that did not bind Omicron RBD (figure 4B). Moreover, these Omicron RBD-binding memory B cells were activated at a similar frequency as non-Omicron RBD-binding memory B cells (figure 4C-D). Although there were no obvious differences in the recruitment of Omicron RBD-binding versus non-binding memory B cells in the recall response to a 3^rd^ dose of wild-type Spike protein, it will be important to determine if heterologous boosting with variant-specific vaccines or variant infection can preferentially recruit cross-reactive or variant-specific memory B cells.

Taken together, our data indicate that mRNA vaccines encoding the original Wuhan Spike generated memory B cells that bind for Omicron and other variant RBDs. These memory B cells were maintained without decline for at least 9-10 months after the primary 2-dose vaccine series. A 3^rd^ dose of mRNA vaccination activated Omicron-specific memory B cells at a similar proportion as Omicron RBD non-binding memory B cells. Thus, mRNA vaccination generates a robust population of memory B cells that maintain reactivity against multiple SARS-CoV-2 VOC, including Omicron, and that are efficiently re-engaged by a 3^rd^ vaccine dose.

### Immune Relationships and Predictors of Boosted Responses

Having quantified antibody and memory B cell responses individually, we next evaluated relationships between different antigen-specific antibody and memory B cell parameters over the course of primary 2-dose vaccination and after a 3^rd^ vaccine dose. To visualize the trajectory of vaccine-induced immunity over time, we clustered samples based on antibody and memory B cell responses using uniform manifold approximation and projection (UMAP) (figure 5A). Infection-naïve and COVID-recovered individuals clustered apart from each other at the pre-vaccination baseline timepoint, as well as at early timepoints following the primary 2-dose vaccine series (figure 5B). Notably, these two groups began to converge in UMAP space at later memory timepoints and were indistinguishable after the 3^rd^ vaccine dose. We also examined correlations between antibody and memory B cell responses over time. Correlation analysis was restricted to individuals without prior COVID or post-vaccine infection. Antibody and memory B cell responses became more tightly correlated at memory timepoints before the 3^rd^ dose (figure 5C, figure S3A), supporting other findings that mRNA vaccines generate coordinated germinal center responses that ultimately result in the export of both long-lived plasma cells and memory B cells (*20*).

**Figure 5.**
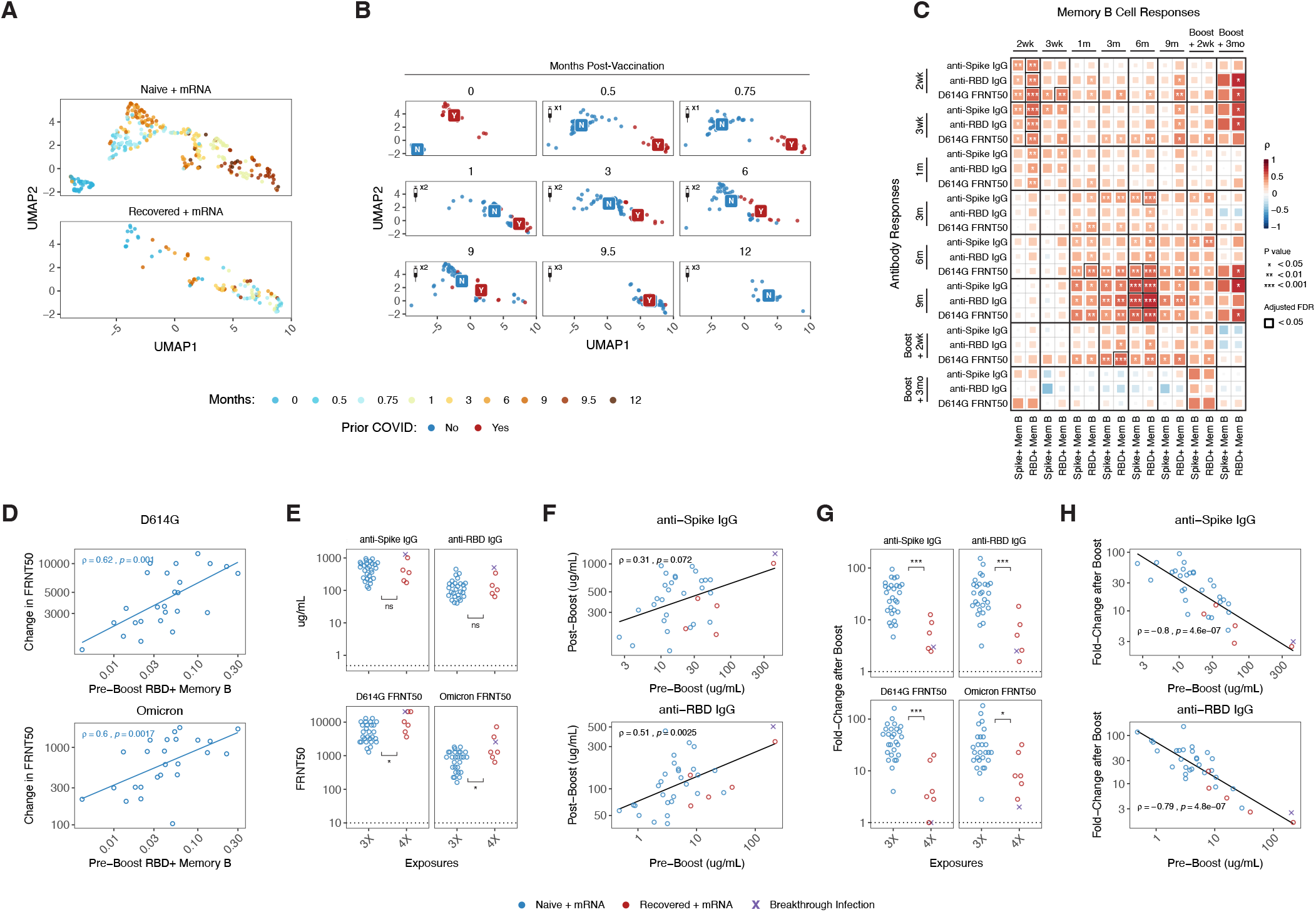
Immune relationships after 2 and 3 doses of mRNA vaccine. **(A)** UMAP of antibody and memory B cell responses to mRNA vaccination. Data points represent individual participants and are colored by timepoint relative to primary vaccine. **B)** UMAP coordinates of SARS-CoV-2– naïve and –recovered subjects over time. Labels indicate centroids for each group at the indicated timepoint. **C)** Correlation matrix of antibody and memory B cell responses over time in SARS-CoV-2–naïve subjects. **D)** Correlation of RBD+ memory B cell frequencies with neutralizing antibody recall responses to D614G and Omicron. Recall responses were calculated as the difference between pre- and post-boost titers ∼2 weeks after the 3^rd^ vaccine dose. **E)** Peak binding and neutralizing antibody responses after 3 versus 4 exposures to SARS-CoV-2 antigen (∼2 weeks post 3^rd^ dose in SARS-CoV-2 naïve and SARS-CoV-2 recovered individuals). **F)** Correlation of pre-boost binding antibody responses with peak post-boost antibody responses. **G)** Fold change in antibody responses after 3 versus 4 exposures to SARS-CoV-2 antigen. **H)** Correlation of fold-change in antibody responses after boosting with pre-3^rd^ dose antibody levels. Statistics were calculated using unpaired non-parametric Wilcoxon test with BH correction for multiple comparisons. All correlations were calculated using nonparametric Spearman rank correlation. *P < 0.05; **P < 0.01; ***P < 0.001; ****P < 0.0001; ns, not significant.

Finally, we investigated how different immune features and additive exposures to SARS-CoV-2 Spike protein affected the absolute and relative magnitude of recall responses. The pre-3^rd^ dose frequency of RBD+ memory B cells correlated with the change in neutralizing antibody titers against D614G and Omicron ∼2 weeks after re-vaccination, consistent with the notion that memory B cells generated after 2 doses of vaccine are an important predictor of subsequent recall responses to SARS-CoV-2 antigens (figure 5D). When comparing individuals with 3 versus 4 total exposures to SARS-CoV-2 Spike antigen, both groups reached similar levels of anti-Spike-and RBD-binding antibodies ∼2 weeks after the 3^rd^ vaccine dose (4^th^ exposure in subjects with previous COVID) (figure 5E). Despite similar binding antibody levels, individuals with 4 total exposures reached slightly higher neutralizing titers against D614G and Omicron (figure 5E). Pre-3^rd^ dose antibody was moderately correlated with peak post-3^rd^ dose antibody levels (figure 5F, figure S3A), but there was considerable variability and pre-3^rd^ dose antibody levels did not fully explain the magnitude of the recall response. To compare the relative benefit of a 3^rd^ vaccine dose, we calculated the fold change in binding and neutralizing antibodies at 2 weeks after re-vaccination compared to paired pre-3^rd^ dose samples. A 3^rd^ vaccine dose induced a ∼10-100-fold increase in antibody titers for individuals with 3 total immune exposures to SARS-CoV-2 Spike (figure 5G). By contrast, the relative recall response to a 3^rd^ vaccine dose in those with previous COVID (i.e., 4 total exposures) was significantly lower with only a ∼5-10 fold-boost (figure 5G).

One concern that has arisen is that vaccinating too soon after a previous exposure might lead to limited boosting. Existing data on this topic for SARS-CoV-2 are so far limited to the time interval between the 1^st^ and 2^nd^ vaccine dose (*31*). Here, we did not observe any significant association between peak antibody titers post-3^rd^ dose and time since primary vaccination (figure S3B; range = 206-372 days), though there was a weak positive association between the fold-change in antibody responses after a 3^rd^ dose and the time since primary vaccination (figure S3C).

Based on these findings, we hypothesized that pre-existing antibody levels may be a key factor that influences the relative magnitude of boosting. Indeed, the pre-3^rd^ dose concentration of anti-Spike or anti-RBD IgG was strongly negatively correlated with the corresponding fold change in antibody after the 3^rd^ vaccine dose (figure 5H). These data suggest a possible feedback mechanism where high levels of circulating antibodies reduce the amount of Spike antigen available to re-activate memory B cells, thereby limiting further antibody production. Accordingly, the relative benefit of boosting with additional vaccine doses will be greatest for individuals who have lower levels of pre-boost antibody.

## Discussion

mRNA vaccination generates protective immunity against SARS-CoV-2 by inducing potent antibody responses as well as memory B cells that can rapidly respond and produce new antibodies upon antigen re-exposure. The establishment of immunity after 2 doses of mRNA vaccine has been well characterized (*15, 29, 32–34*). However, it remains unclear how additional vaccine doses and combinations of vaccination and infection affect the quantity and quality of immune responses, particularly against immune-evasive SARS-CoV-2 variants like Omicron. In this study, we examined antibody and memory B cell responses through ∼9 months following the primary 2-dose vaccine series as well as up to ∼3 months following a 3^rd^ vaccine dose. In particular, the longitudinal nature of this study enabled detailed analysis of the magnitude, durability, and quality of SARS-CoV-2 vaccine-induced immunity over ∼1 year and multiple antigen exposures.

This study provides several key pieces of data relevant to SARS-CoV-2 and mRNA vaccine immunobiology. Although antibody levels declined from peak levels ∼1 week after the second dose to ∼6 months post-primary vaccination, antibody titers then stabilized between 6- and 9-months post-vaccination with continued improvement of neutralization potency over this period. A 3^rd^ vaccine dose at ∼9 months post-primary vaccination increased antibody responses ∼10 to 100-fold, including boosting neutralizing antibodies against the Omicron variant. Moreover, the antibody levels achieved after the 3^rd^ dose were similar to those observed in SARS-CoV-2 recovered individuals after 2 doses of mRNA vaccine (i.e. hybrid immunity). Breakthrough infection after 2 doses of mRNA vaccine also appeared to produce similar antibody boosting as a 3^rd^ vaccine dose. These boosted antibody responses subsequently declined over time but still remained significantly above pre-boost levels at 3 months post 3^rd^ dose. In contrast to antibodies, that decayed over time following vaccination, memory B cells numbers remained highly stable in the blood with no evidence of decay at ∼9 months post-primary vaccination. Notably, 2-dose vaccination generated a robust memory B cell response against the Omicron variant, with ∼40-50% of RBD-binding memory B cells able to cross-bind Alpha, Beta, Delta, and Omicron. A 3^rd^ vaccine dose efficiently recruited memory cells with cross-reactivity to multiple VOCs, resulting in amplification of antibody responses capable of neutralizing heterologous Spike proteins from immune-evasive SARS-CoV-2 VOCs including Omicron. The ability of approximately half of the memory B cell pool to bind multiple variants indicates that the antibodies encoded by these memory B cells are targeting more conserved epitopes of RBD or are of higher quality and able to overcome epitope changes associated with mutations in VOCs. It will be important to determine how resilient these memory B cell responses are to emerging variants. For example, BA.2 shares many features with previous VOCs but also contains additional mutations not observed in the BA.1 Omicron sublineage (*35*). Regardless, mRNA vaccines containing the original Wuhan Spike appear to be capable of generating and boosting memory B cell responses and associated antibodies with the capacity to recognize all current SARS-CoV-2 VOCs and may provide lasting protection against future variants.

Our data also identify several factors that predict the absolute and relative benefit of boosting. Binding antibody levels achieved after a 3^rd^ dose were similar in individuals with or without a prior SARS-CoV-2 infection. Despite similar anti-Spike and anti-RBD antibody titers, infected then vaccinated individuals with 4 total immune exposures reached slightly higher neutralizing titers against D614G and Omicron than SARS-CoV-2 naïve individuals who received 3 doses of vaccine, suggesting that additional vaccination or infection following a 3-dose vaccine regimen still has a quantitative benefit to antibody responses. Although boosting with a 3^rd^ vaccine dose universally increased the total magnitude of binding and neutralizing antibody responses compared to pre-boost levels, a 3^rd^ exposure to SARS-CoV-2 Spike antigen increased antibody levels 10- to 100-fold, whereas a 4^th^ exposure in individuals with higher pre-boost antibodies resulted in less robust boosting (∼5- to 10-fold). A major factor influencing the magnitude of the boost was pre-3^rd^ vaccine dose antibody concentrations, which were inversely correlated with the fold-change of antibody boosting. These observations suggest that pre-existing high titers of antibody may compete with memory B cells for antigen and limit the extent of antibody boosting. Accordingly, the relative value of additional vaccine doses will likely be greatest for individuals with lower pre-boost antibodies, including immunocompromised or older populations.

Overall, these data support the utility of a 3^rd^ vaccine dose, but also highlight immunological constraints that may set a limit on maximum antibody levels achieved by repeated short interval boosting. Additional work is required to evaluate variant-specific vaccines and how boosting with a modified antigen may augment recall responses compared to boosting with the original Wuhan strain sequence (*36*). Nevertheless, these data highlight the urgent need to better understand what antibody titers are necessary for protection against infection and/or severe disease (*28*). Should this threshold be defined, it may be useful to implement serologic testing to maximize the benefit and equity of additional vaccine doses moving forward.

## Acknowledgements

We thank the study participants for their generosity in making the study possible. We also thank Ali Ellebedy and Julian Zhou for helpful discussions and feedback, as well as the Flow Cytometry Core at the University of Pennsylvania for technical support.

## Methods

### Clinical Recruitment and Sample Collection

61 individuals (45 SARS-CoV-2 naïve, 16 SARS-CoV-2 recovered) were consented and enrolled in the longitudinal vaccine study with approval from the University of Pennsylvania Institutional Review Board (IRB# 844642). All participants were otherwise healthy and based on self-reported health screening did not have any history of chronic health conditions. Subjects were stratified based on self-reported and laboratory evidence of a prior SARS-CoV-2 infection. All subjects received either Pfizer (BNT162b2) or Moderna (mRNA-1273) mRNA vaccines. Samples were collected at 10 timepoints: baseline, 2 weeks post-1^st^ dose, day of 2^nd^ dose, 1 week post-2^nd^ dose, 3 months post-primary immunization, 6 months post-primary immunization, 9-10 months post-primary immunization, pre 3^rd^ dose, 2 weeks post 3^rd^ dose, and 3 months post 3^rd^ dose. 80-100mL of peripheral blood samples and clinical questionnaire data were collected at each study visit. Full cohort and demographic information is provided in table S1. Healthy donor PBMC samples were collected with approval from the University of Pennsylvania Institutional Review Board (IRB# 845061)

### Peripheral Blood Sample Processing

Venous blood was collected into sodium heparin and EDTA tubes by standard phlebotomy. Blood tubes were centrifuged at 3000rpm for 15 minutes to separate plasma. Heparin and EDTA plasma were stored at −80°C for downstream antibody analysis. Remaining whole blood was diluted 1:1 with RPMI + 1% FBS + 2mM L-Glutamine + 100 U Penicillin/Streptomycin and layered onto SEPMATE tubes (STEMCELL Technologies) containing lymphoprep gradient (STEMCELL Technologies). SEPMATE tubes were centrifuged at 1200g for 10 minutes and the PBMC fraction was collected into new tubes. PBMCs were then washed with RPMI + 1% FBS + 2mM L-Glutamine + 100 U Penicillin/Streptomycin and treated with ACK lysis buffer (Thermo Fisher) for 5 minutes. Samples were washed again with RPMI + 1% FBS + 2mM L-Glutamine + 100 U Penicillin/Streptomycin, filtered with a 70µm filter, and counted using a Countess automated cell counter (Thermo Fisher). Aliquots containing 5-10×10^6^ PBMCs were cryopreserved in fresh 90% FBS 10% DMSO.

### Detection of SARS-CoV-2 Spike- and RBD-Specific Antibodies

Plasma samples were tested for SARS-CoV-2-specific antibody by enzyme-linked immunosorbent assay (ELISA) as described (*15, 18, 37*). Plasmids encoding the recombinant full-length Spike protein and the RBD were provided by F. Krammer (Mt. Sinai) and purified by nickel-nitrilotriacetic acid resin (Qiagen). ELISA plates (Immulon 4 HBX, Thermo Fisher Scientific) were coated with PBS or 2 ug/mL recombinant protein and stored overnight at 4C. The next day, plates were washed with phosphate-buffered saline containing 0.1% Tween-20 (PBS-T) and blocked for 1 hour with PBS-T supplemented with 3% non-fat milk powder. Samples were heat-inactivated for 1 hour at 56C and diluted in PBS-T supplemented with 1% non-fat milk powder. After washing the plates with PBS-T, 50 uL diluted sample was added to each well. Plates were incubated for 2 hours and washed with PBS-T. Next, 50 uL of 1:5000 diluted goat anti-human IgG-HRP (Jackson ImmunoResearch Laboratories) or 1:1000 diluted goat anti-human IgM-HRP (SouthernBiotech) was added to each well and plates were incubated for 1 hour. Plates were washed with PBS-T before 50 uL SureBlue 3,3’,5,5’-tetramethylbenzidine substrate (KPL) was added to each well. After 5 minutes incubation, 25 uL of 250 mM hydrochloric acid was added to each well to stop the reaction. Plates were read with the SpectraMax 190 microplate reader (Molecular Devices) at an optical density (OD) of 450 nm. Monoclonal antibody CR3022 was included on each plate to convert OD values into relative antibody concentrations. Plasmids to express CR3022 were provided by I. Wilson (Scripps).

### Detection of SARS-CoV-2 Neutralizing Antibodies

293T cells were seeded for 24 hours at 5 × 10^6^ cells per 10 cm dish and were transfected using calcium phosphate with 25 μg of pCG1 SARS-CoV-2 S D614G delta18, pCG1 SARS-CoV-2 S B.1.617.2 delta 18, or pCG1 SARS-CoV-2 S B.1.1.529 delta 18 expression plasmid encoding a codon optimized SARS-CoV-2 S gene with an 18-residue truncation in the cytoplasmic tail. Variant sequences are provided below. 12 hours post transfection, cells were fed with fresh media containing 5mM sodium butyrate to increase expression of the transfected DNA. 24 hours after transfection, the SARS-CoV-2 Spike expressing cells were infected for 2 hours with VSV-G pseudotyped VSVΔG-RFP at an MOI of ∼1-3. Virus containing media was removed and the cells were re-fed with media without serum. Media containing the VSVΔG-RFP SARS-CoV-2 pseudotypes was harvested 28-30 hours after infection, clarified by centrifugation twice at 6000g, then aliquoted and stored at −80 °C until used for antibody neutralization analysis. All sera were heat-inactivated for 30 minutes at 53 ⁰C prior to use in the neutralization assay. Vero E6 cells stably expressing TMPRSS2 were seeded in 100 μl at 2.5×10^4^ cells/well in a 96 well collagen coated plate. The next day, 2-fold serially diluted serum samples were mixed with VSVΔG-RFP SARS-CoV-2 pseudotype virus (100-300 focus forming units/well) and incubated for 1 hour at 37⁰C. 1E9F9, a mouse anti-VSV Indiana G, was also included in this mixture at a concentration of 600 ng/ml (Absolute Antibody, Ab01402-2.0) to neutralize any potential VSV-G carryover virus. The serum-virus mixture was then used to replace the media on VeroE6 TMPRSS2 cells. 21-22 hours post-infection, the cells were washed and fixed with 4% paraformaldehyde before visualization on an S6 FluoroSpot Analyzer (CTL, Shaker Heights OH). Individual infected foci were enumerated and the values were compared to control wells without antibody. The focus reduction neutralization titer 50% (FRNT_50_) was measured as the greatest serum dilution at which focus count was reduced by at least 50% relative to control cells that were infected with pseudotype virus in the absence of human serum. FRNT_50_ titers for each sample were measured in at least two technical replicates and were reported for each sample as the geometric mean of the technical replicates.

| Variant | Mutations in Pseudovirus Construct |
| --- | --- |
| D614G (WT) | D614G |
| B.1.351 (Beta) | L18F, D80A, D215G, R246I, K417N, E484K, N501Y, D614G, A701V |
| B.1.617.2 (Delta) | T19R, G142D, del156-157, R158G, L452R, T478K, D614G, P681R, D950N |
| B.1.1.529 (Omicron) | A67V, del69-70, T95I, G142D, del143-145, N211I, del212, ins214 (EPE), G339D, S371L, S373P, S375F, S477N, T478K, E484A, Q493R, G496S, Q498R, N501Y, Y505H, T547K, D614G, G655Y, N679K, P681H, D796Y, N856K, Q954H, N969K, L981F |

### Detection and Phenotyping of SARS-CoV-2-Specific Memory B Cells

Antigen-specific B cells were detected using biotinylated proteins in combination with different streptavidin (SA)-fluorophore conjugates as described (*15, 18*). All reagents are listed in table S2. Biotinylated proteins were multimerized with fluorescently labeled SA for 1 hour at 4C. Full-length Spike protein was mixed with SA-BV421 at a 10:1 mass ratio (200ng Spike with 20ng SA; ∼4:1 molar ratio). Spike RBD was mixed with SA-APC at a 2:1 mass ratio (25ng RBD with 12.5ng SA; ∼4:1 molar ratio). Biotinylated influenza HA pools were mixed with SA-PE at a 6.25:1 mass ratio (100ng HA pool with 16ng SA; ∼6:1 molar ratio). Influenza HA antigens corresponding with the 2019 trivalent vaccine (A/Brisbane/02/2018/H1N1, B/Colorado/06/2017) were chosen as a historical antigen and were biotinylated using an EZ-Link Micro NHS-PEG4 Biotinylation Kit (Thermo Fisher) according to the manufacturer’s instructions. Excess biotin was subsequently removed using Zebra Spin Desalting Columns 7K MWCO (Thermo Fisher) and protein was quantified with a Pierce BCA Assay (Thermo Fisher). SA-BV711 was used as a decoy probe without biotinylated protein to gate out cells that non-specifically bind streptavidin. All experimental steps were performed in a 50/50 mixture of PBS + 2% FBS and Brilliant Buffer (BD Bioscience). Antigen probes for Spike, RBD, and HA were prepared individually and mixed together after multimerization with 5uM free D-biotin (Avidity LLC) to minimize potential cross-reactivity between probes. For staining, 5×10^6^ cryopreserved PBMC samples were prepared in a 96-well U-bottom plate. Cells were first stained with Fc block (Biolegend, 1:200) and Ghost 510 Viability Dye for 15 minutes at 4C. Cells were then washed and stained with 50uL antigen probe master mix containing 200ng Spike-BV421, 25ng RBD-APC, 100ng HA-PE, and 20ng SA-BV711 decoy for 1 hour at 4C. Following incubation with antigen probe, cells were washed again and stained with anti-CD3, anti-CD19, anti-CD20, anti-CD27, anti-CD38, anti-CD71, anti-IgD, anti-IgM, anti-IgG, and anti-IgA for 30 minutes at 4C. After surface stain, cells were washed and fixed in 1% PFA overnight at 4C. Antigen-specific gates for B cell probe assays were set based on healthy donors stained without antigen probes (similar to an FMO control) and were kept the same for all experimental runs.

### Detection of Variant RBD, NTD, and S2-Specific Memory B Cells

Variant RBD, NTD, and S2-specific memory B cells were detected using a similar approach as described above. SARS-CoV-2 nucleocapsid was used as a vaccine-irrelevant antigen control. All reagents are listed in table S2. Probes were multimerized for 1.5 hours at the following ratios (all ∼4:1 molar ratios calculated relative to the streptavidin-only component irrespective of fluorophore): 200ng full-length Spike protein was mixed with 20ng SA-BV421, 30ng N-terminal domain was mixed with 12ng SA-BV786, 25ng wild-type RBD was mixed with 12.5ng SA-BB515, 25ng Alpha RBD was mixed with 12.5ng SA-BV711, 25ng Beta RBD was mixed with 12.5ng SA-PE, 25ng Delta was mixed with 12.5ng SA-APC, 25ng Omicron was mixed with 12.5ng SA-PE-Cy7, 50ng S2 was mixed with 12ng SA-BUV737, 50ng nucleocapsid was mixed with 14ng SA-BV605. 12.5ng SA-BUV615 was used as a decoy probe. All antigen probes were multimerized separately and mixed together with 5uM free D-biotin. Prior to staining, total B cells were enriched from 10-20×10^6^ cryopreserved PBMC samples by negative selection using an EasySep human B cell isolation kit (STEMCELL, #17954). B cells were then prepared in a 96-well U-bottom plate and stained with Fc block and Ghost 510 Viability Dye as described above. Cells were washed and stained with 50uL antigen probe master mix for 1 hour at 4C. After probe staining, cells were washed again and stained with anti-CD3, anti-CD11c, anti-CD19, anti-CD21, anti-CD27, anti-CD38, and anti-IgD for 30 minutes at 4C. After surface stain, cells were washed and fixed in 1X Stabilizing Fixative (BD Biosciences) overnight at 4C.

### Flow Cytometry

Samples were acquired on a BD Symphony A5 instrument. Standardized SPHERO rainbow beads (Spherotech) were used to track and adjust photomultiplier tubes over time. UltraComp eBeads (Thermo Fisher) were used for compensation. Up to 5×10^6^ cells were acquired per sample. Data were analyzed using FlowJo v10 (BD Bioscience). For Boolean analysis of variant cross-binding, data were imported into SPICE 6 (NIH Vaccine Research Center (*38*)).

### High Dimensional Analysis and Statistics

All data were analyzed using custom scripts in R and visualized using RStudio. Pairwise correlations between variables were calculated and visualized as a correlogram using corrplot with FDR correction as described previously (*39*). For heatmaps, data were visualized with pheatmap. For construction of UMAPs, 5 antigen-specific immune features were selected: anti-Spike IgG, anti-RBD IgG, D614G FRNT50, Spike+ memory B cells, and RBD+ memory B cells. Antibody and cell frequency data were log10 transformed and scaled by column (z-score normalization) before generating UMAP coordinates. Statistical tests are indicated in the corresponding figure legends. All tests were performed two-sided with a nominal significance threshold of p < 0.05. Benjamini-Hochberg (BH) correction was performed in all cases of multiple comparisons. Unpaired tests were used for comparisons between timepoints unless otherwise indicated as some participants were missing samples from individual timepoints. * indicates p < 0.05, ** indicates p < 0.01, *** indicates p < 0.001, **** indicates p < 0.0001. Source code and data files are available upon request from the authors.

## Supplemental Figures

**Figure S1.**
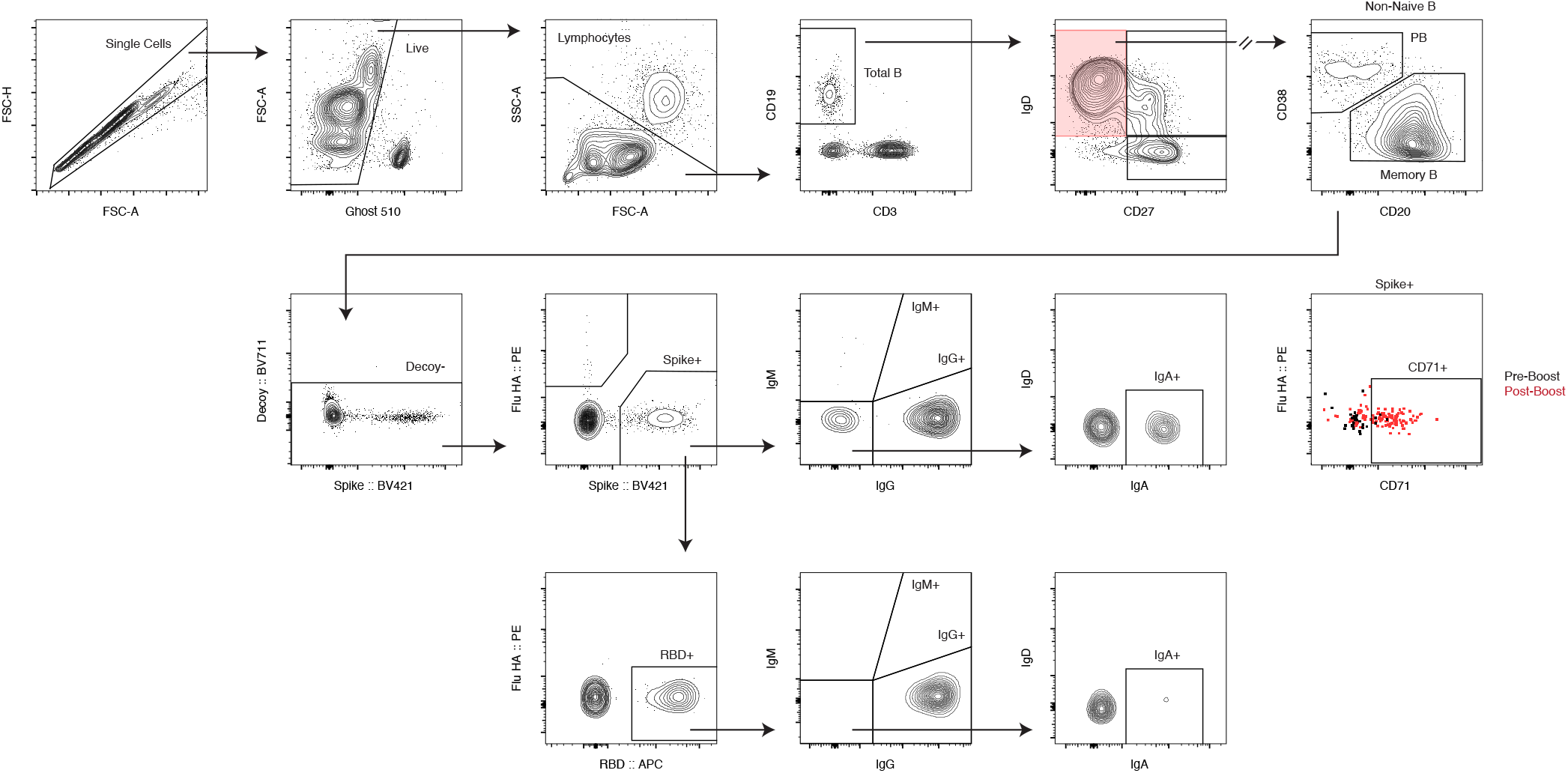
Gating strategy for identifying SARS-CoV-2-specific B cell responses. Single cells were identified based on FSC-A and FSC-H. Live cells were identified based on negative staining with Ghost 510 viability dye. Lymphocytes were identified from bulk PBMCs based on FSC-A and SSC-A. Total B cells were then identified from live lymphocytes as CD3-CD19+ cells. Memory cell subsets were identified based on expression of IgD, CD20, CD27, and CD38. IgD+ CD27-naïve cells were excluded from all analysis. From non-naïve B cells, memory B cells were identified as CD20+ CD38lo/intermediate and plasmablasts were identified as CD20lo CD38++. Antigen-specificity of memory cells and plasmablasts was determined based on binding to fluorescently labelled antigen probes. First, a decoy probe (BV711-streptavidin) was used to exclude cells that non-specifically bind streptavidin. Decoy-cells were then assessed for binding to full-length SARS-CoV-2 Spike protein or influenza hemagglutinin (HA) from the 2019 flu vaccine season. Spike+ cells were subsequently analyzed for co-binding to a receptor-binding domain (RBD) probe. Regarding phenotype, immunoglobulin isotype (IgG, IgM, IgA) was measured on all antigen-binding populations. CD71 was used as a marker of activated B cells.

**Figure S2.**
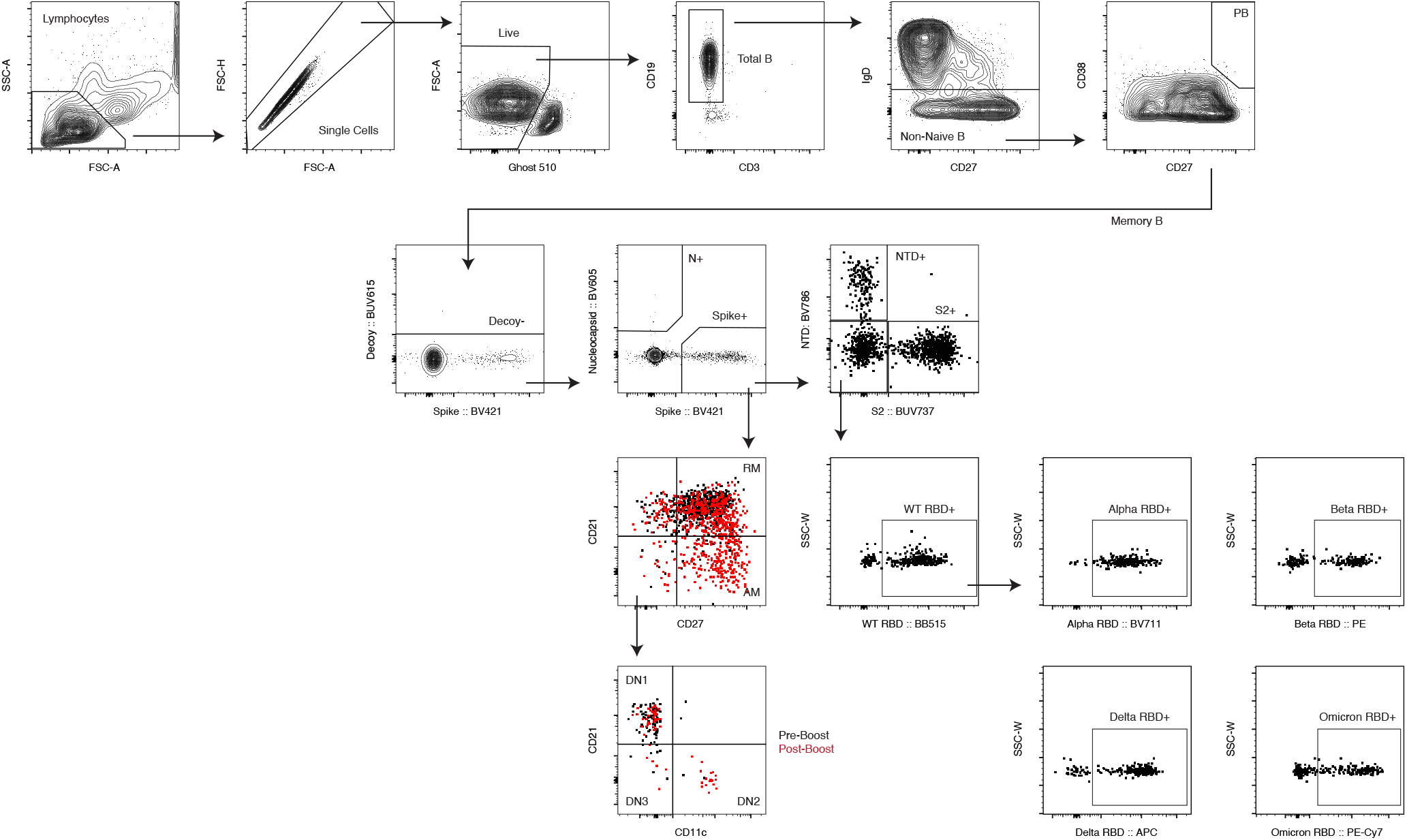
Gating strategy for identifying SARS-CoV-2 variant-reactive memory B cell responses. B cells were pre-enriched from total PBMCs by negative selection with a STEMCELL isolation kit. Lymphocytes were identified based on FSC-A and SSC-A. Single cells were identified based on FSC-A and FSC-H. Live cells were identified based on negative staining with Ghost 510 viability dye. Total B cells were then identified from live lymphocytes as CD3-CD19+ cells. IgD+ CD27-naïve cells were excluded from all analysis. From non-naïve B cells, memory B cells were identified as CD20+ CD38lo/intermediate. Antigen-specificity of memory cells and plasmablasts was determined based on binding to fluorescently labelled antigen probes. First, a decoy probe (BUV615-streptavidin) was used to exclude cells that non-specifically bind streptavidin. Decoy-cells were then assessed for binding to full-length SARS-CoV-2 Spike protein or SARS-CoV-2 Nucleocapsid protein. Spike+ cells were subsequently analyzed for co-binding to NTD and S2 probes. Spike+ cells that did not bind NTD or S2 were analyzed for binding to wild-type receptor-binding domain (RBD) probe. Cross-binding to Alpha, Beta, Delta, and Omicron variant RBD probes was measured on WT RBD-specific cells. Regarding phenotype, memory subsets were identified based on CD11c, CD21, and CD27 staining. CD21+ CD27+ cells were defined as resting memory and CD21-CD27+ cells were defined as activated memory. CD27-memory cells were split into double negative (DN) subsets based on CD11c and CD21 staining.

**Figure S3.**
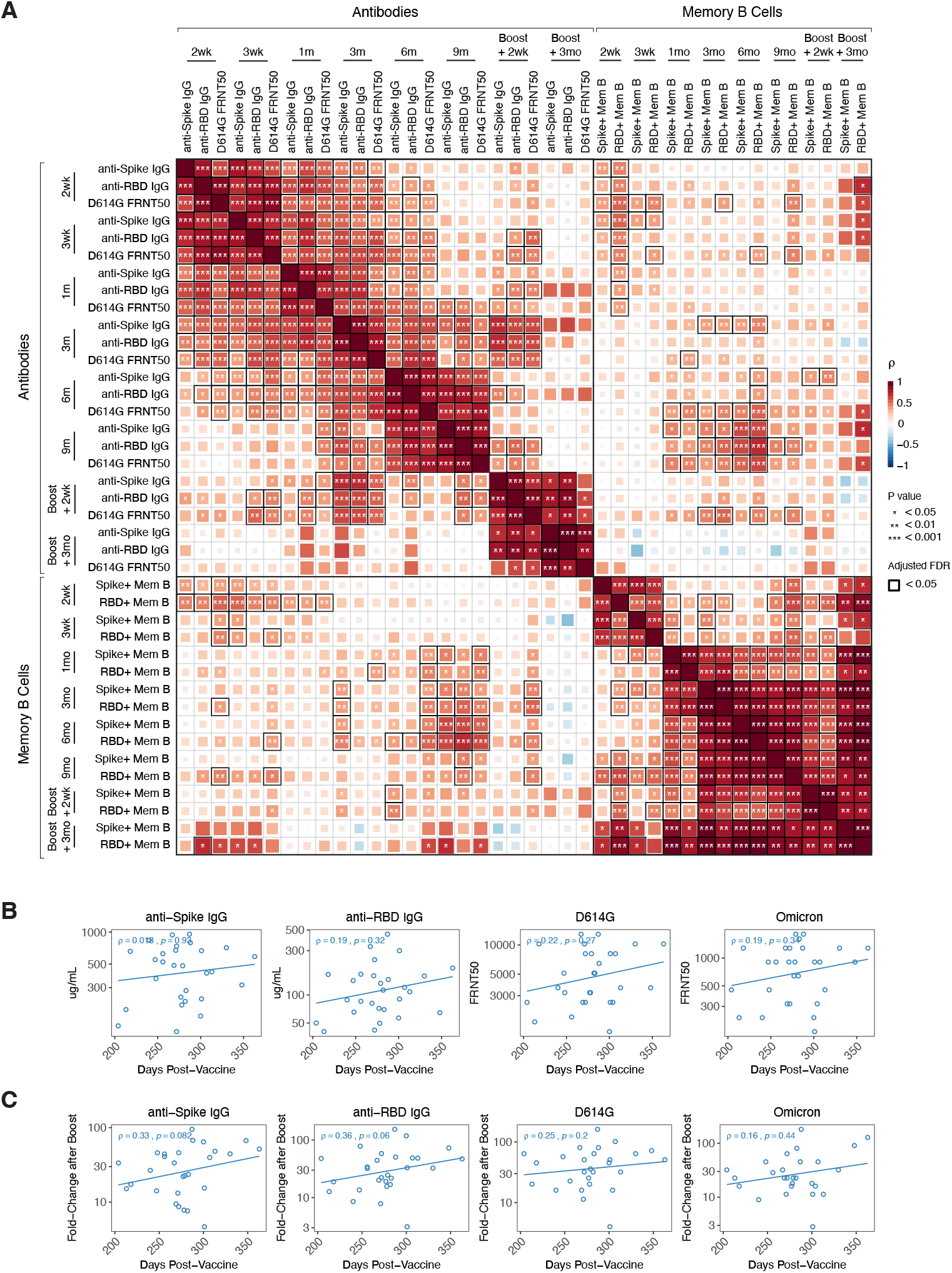
Immune relationships and correlation of antibody boosting with time since primary vaccination. **A)** Correlation matrix of antibody and memory B cell responses over time in SARS-CoV-2–naïve subjects. **B)** Correlation of peak post-boost antibody levels (∼2 weeks after the 3^rd^ dose) with days since primary vaccination. **C)** Correlation of fold-change in antibody responses after boosting with days since primary vaccination. All correlations were calculated using nonparametric Spearman rank correlation. *P < 0.05; **P < 0.01; ***P < 0.001; ****P < 0.0001; ns, not significant.

**Table S1.** Clinical metadata.

| Characteristics |  | SARS-CoV-2 Naïve | SARS-CoV-2 Recovered |
| --- | --- | --- | --- |
| Total | Number | 45 | 16 |
| Age | Average (Years) | 36.9 | 38.3 |
|  | Range (Years) | 22-67 | 23-59 |
|  | 20-30 | 15 | 4 |
|  | 30-40 | 14 | 6 |
|  | 40-50 | 9 | 2 |
|  | 50+ | 7 | 4 |
| Sex | Male | 21 | 10 |
|  | Female | 24 | 6 |
| Race/Ethnicity | White - Non-Hispanic/Latino | 27 | 7 |
|  | White - Hispanic/Latino | 4 | 1 |
|  | Asian | 8 | 6 |
|  | Black | 2 | 1 |
|  | Native | 0 | 1 |
|  | Other | 1 | 0 |
| Primary Vaccine Type | Pfizer | 42 | 12 |
|  | Moderna | 3 | 4 |
| Time Between Infection and Vaccine | Average (Days) | -- | 102.4 |
|  | Range (Days) | -- | 50-275 |
| Booster (3<sup>rd</sup>) Vaccination | Yes | 37 | 10 |
|  | Average Time Relative to 1<sup>st</sup> Dose (Days) | 280.4 | 308.6 |
|  | Range (Days) | 206-372 | 277-366 |
| Breakthrough Infection | After 2<sup>nd</sup> Dose | 3 | 1 |
|  | After 3<sup>rd</sup> Dose | 4 | 1 |

**Table S2.**
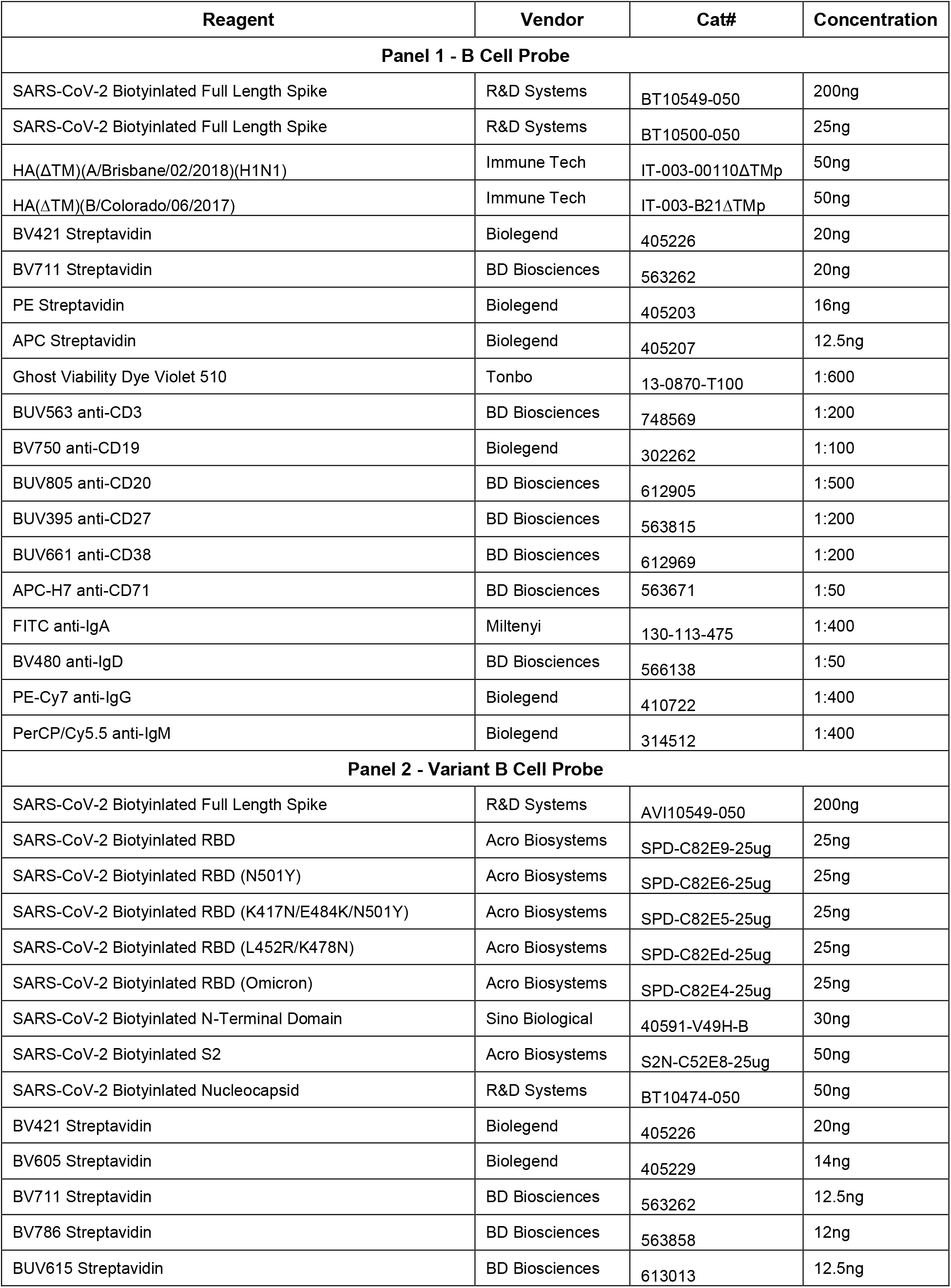
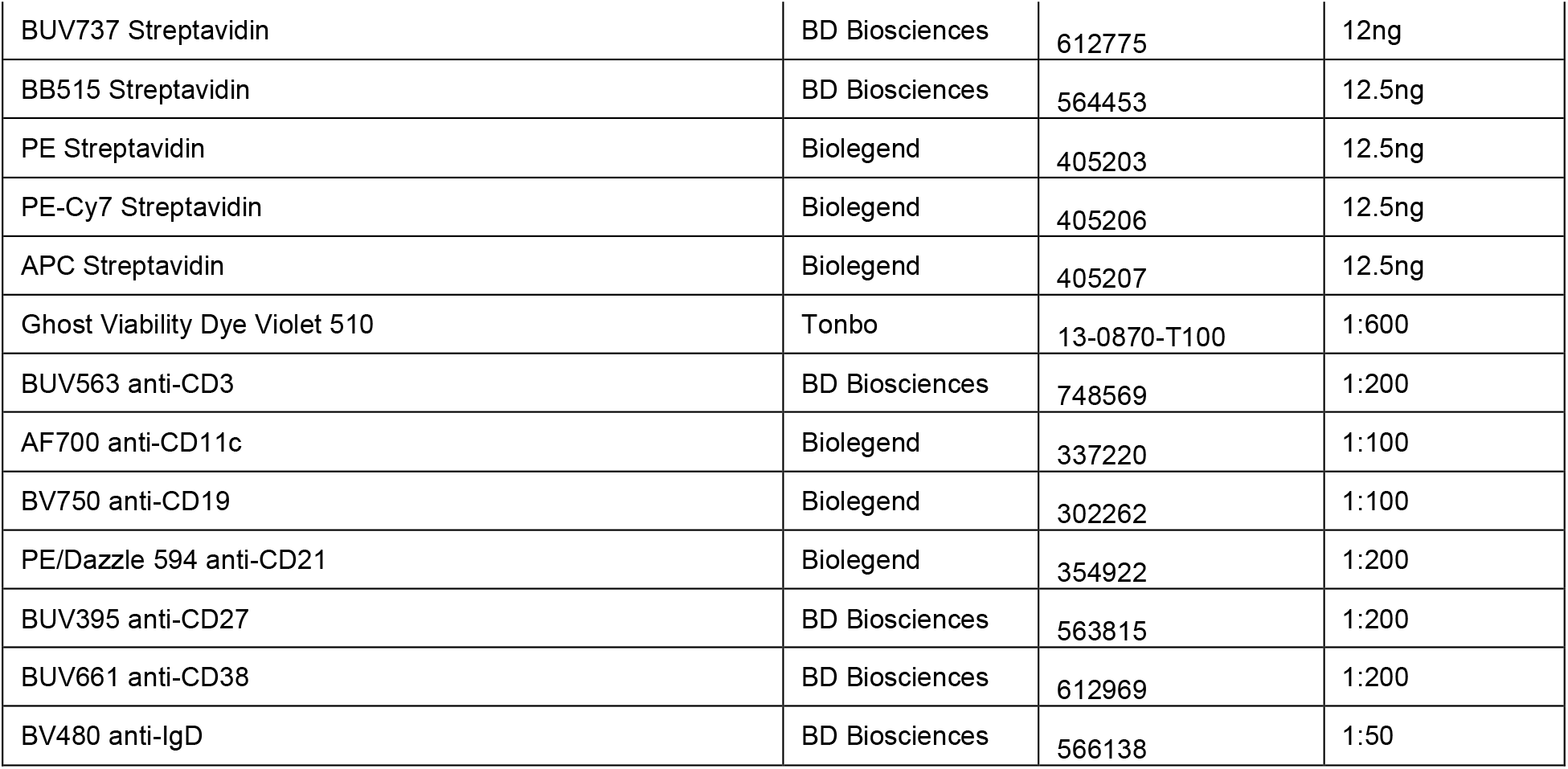
Reagent information.

